# Distinct Laminar Processing of Local and Global Context in Primate Primary Visual Cortex

**DOI:** 10.1101/171793

**Authors:** Maryam Bijanzadeh, Lauri Nurminen, Sam Merlin, Alessandra Angelucci

**Author notes:** Corresponding author address: 65 Mario Capecchi Drive, Salt Lake City, UT 84132, USA. Present address: Department of Neurological Surgery, UCSF, CA 94143, USA. Present address: Medical Science, School of Science and Health, Western Sydney University, Campbelltown, NSW 2560 Australia.

## Abstract

Visual perception is profoundly affected by spatial context. In visual cortex, neuronal responses to stimuli inside their receptive field (RF) are suppressed by contextual stimuli in the RF surround (surround suppression). How do neuronal RFs integrate information across visual space, and what role do different cortical layers play in the processing of spatial context? By recording simultaneously across all layers of macaque primary visual cortex, while presenting visual stimuli at increasing distances from the recorded cells’ RF, we find that near vs. far stimuli activate distinct layers. Stimuli in the near-surround evoke the earliest subthreshold responses in superficial and deep layers, and cause the earliest surround suppression of spiking responses in superficial layers. Instead, far-surround stimuli evoke the earliest subthreshold responses in feedback-recipient layers, i.e. 1 and the lower half of the deep layers, and suppress visually-evoked spiking responses almost simultaneously in all layers, except 4C, where suppression emerges latest. Our results reveal unique contributions of the cortical layers to the processing of local and global spatial context, and suggest distinct underlying circuits for local and global signal integration.

## I. INTRODUCTION

The mammalian neocortex consists of six interconnected layers with distinct functional properties and input/output connections. This laminar architecture is a prominent feature of all neocortex, yet its role in information processing remains elusive. To understand laminar processing, we have chosen to study a canonical cortical computation found across sensory modalities and species, surround suppression (reviewed in: Allman et al., 1985; Angelucci and Shushruth, 2013), in a cortical area whose laminar connectivity and neuronal response properties are well understood, i.e. the macaque primary visual cortex (V1) (Callaway, 2014; Hubel and Wiesel, 2004). Understanding the role of V1 layers in surround suppression has the potential to reveal fundamental principles of laminar computation across sensory systems.

Surround suppression is a form of contextual modulation, which endows neurons with the ability to change their responses to stimuli inside their receptive field (RF) depending on spatial context, i.e. the stimuli outside the RF (Angelucci and Shushruth, 2013; Blakemore and Tobin, 1972; Hubel and Wiesel, 1965). These modulatory effects are typically suppressive when the stimuli in the RF and surround have similar properties. In the visual cortex, surround suppression is thought to contribute to important perceptual phenomena, such as segmentation of object boundaries and visual saliency (Knierim and Van Essen, 1992; Nothdurft et al., 2000; Nurminen and Angelucci, 2014; Petrov and McKee, 2006).

Traditional single-unit electrophysiology has revealed laminar differences in the properties of surround suppression (Henry et al., 2013; Ichida et al., 2007; Sceniak et al., 2001; Shushruth et al., 2009, 2013). However, it remains unknown how neurons in different cortical layers integrate visual signals from outside their RFs, and in which layers surround suppression first emerges. Moreover, because cortical layers exhibit different patterns of afferent connections, understanding which layers integrate signals from the RF surround and generate surround suppression can reveal the circuitry underlying surround phenomena and contextual integration. Specifically, in macaque V1, feedforward afferents from the lateral geniculate nucleus (LGN) terminate primarily in layer 4C (Blasdel and Lund, 1983; Hubel and Wiesel, 1972); millimeters-long horizontal connections are present in all V1 layers, except lower 4C, but are most prominent in layers 2/3 and 5, and weaker in 4B-upper 4Cα and 6 (Angelucci et al., 2002a; Lund et al., 2003; Rockland and Lund, 1983); feedback afferents from higher visual areas terminate primarily in V1 layers 1-2A and 5B-6 (Federer et al., 2015; Rockland and Pandya, 1979) (**Fig. 1**). Given this laminar specificity of afferents to V1, we reasoned that the specific circuits carrying surround signals to a V1 column evoke the earliest postsynaptic depolarization in the V1 layers where they terminate.

**Figure 1.**
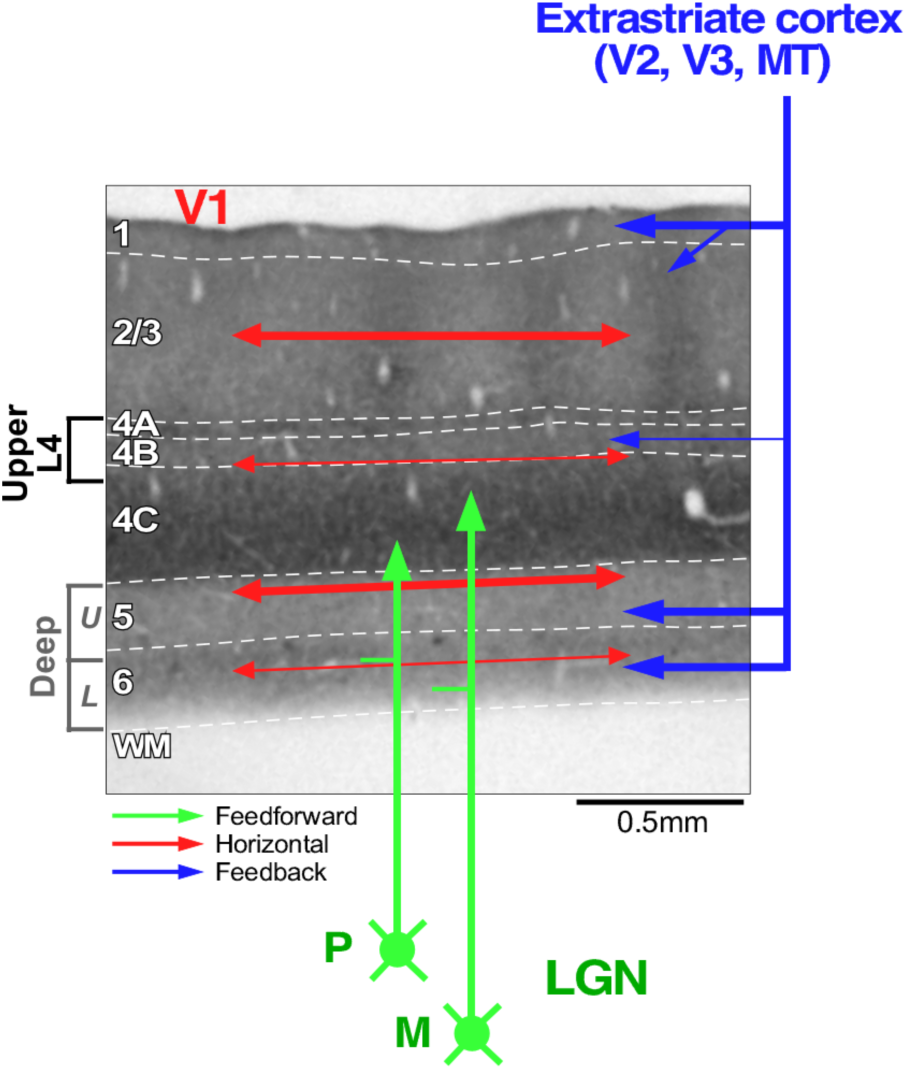
Laminar Specificity of Feedforward, Horizontal and Feedback Projections to V1. (A) Schematics of V1 laminar terminations of geniculocortical (*green arrows*), intra-V1 horizontal (*red arrows*) and inter-areal feedback (*blue arrows*) projections to V1, shown on a pia-to-white matter (*WM*) section of macaque V1 stained for the metabolic enzyme cytochrome oxidase to reveal layers. Arrow thickness represents density of projections. *White dashed contours:* laminar boundaries; layers are indicated to the left. The terminology used in this study to indicate layers above and below L4C is shown to the left of the image; specifically, *Deep_U_*, *Deep_L_* indicate the upper and lower half of the deep layers, respectively, and *Upper*-*L4* encompasses upper 4Cα, L4B and L4A. *M*, *P:* magnocellular and parvocellular LGN inputs, respectively.

To understand the contribution of V1 layers and of different connection types to surround suppression, we recorded simultaneously through all layers of macaque V1 the local field potential (LFP) and multiunit spiking activity (MUA) evoked by visual stimuli presented at increasing distances from the recorded neurons RF, and measured the onset latency of subthreshold responses and of surround suppression through the layers. Our results reveal that distinct layers, and therefore distinct circuits, are involved in the processing of local and global spatial context.

## II. RESULTS

We recorded visually-evoked LFP and MUA using 24 channel linear electrode arrays (100 μm electrode spacing) oriented perpendicular to the cortical surface of area V1 in sufentanil-anesthetized and paralyzed macaque monkeys (see Methods). The verticality of the array was verified *in vivo* by the alignment of receptive field (RF) location and similarity of orientation tuning functions through the depth of the cortex (**Fig. 2A**), as well as by postmortem histology (**Fig. 2B**). Here we present results from neuronal activity recorded in 4 macaques at 162 contacts from 10 penetrations that were deemed to be perpendicular to the V1 surface by these criteria.

**Figure 2.**
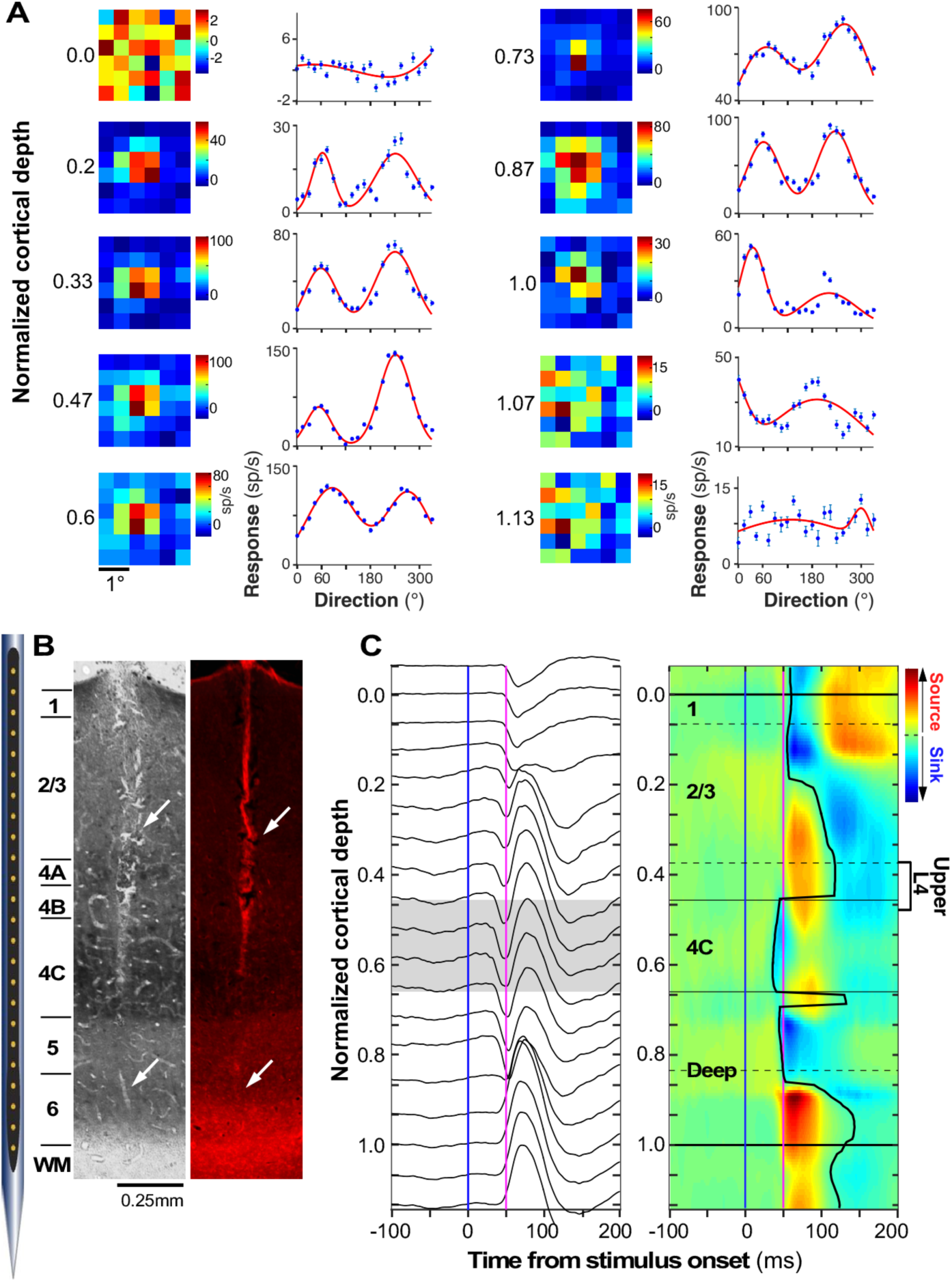
Identification of Laminar Boundaries and RF Alignment in a V1 Column. **(A) <u>Left columns</u>:** Minimum response field (mRF) mapping shown as a color map for every other contact through the depth of the cortex, obtained by averaging the MUA response (0-200ms after stimulus onset) to 0.5° black square stimuli flashed in a 6×6 grid centered on the mRF. Note that the mRFs (hot spots) across contacts are well aligned, indicating perpendicular penetration. **<u>Right columns</u>:** direction tuning curves obtained at each contact in response to 1° diameter grating patches of varying orientation and direction centered on the aggregate mRF of the column. Similarity of orientation preference across contacts indicates perpendicular penetration. *Red curves:* fits to the data (*blue dots*) using the sum of two Gaussians. Numbers to the left of the mRF maps indicate normalized cortical depth, with 0 and 1.0 indicating top and bottom of the cortex, respectively. **(B) <u>Left</u>**: Cytochrome oxidase staining of pia-to-white matter section of V1 showing the array track as a discoloration in staining (*arrows*). **<u>Right</u>:** Same section viewed under fluorescence showing DiI staining of the array track (arrows). **(C) <u>Left</u>:** Stimulus-evoked LFP profile from same array penetration as in (A-B) obtained in response to a flashing 0.5° black square centered on the mRF of the V1 column. *Gray shading:* L4C. **<u>Right</u>:** Baseline-corrected (z-scored; see Methods) CSD calculated from the LFPs and displayed as a color map. *Black contour* on the CSD represents the onset latency of current sinks. *Solid and dashed horizontal lines* mark the main cortical boundaries and their subdivisions, respectively. *Blue and purple vertical bars* on the LFP and CSD profiles mark the stimulus onset (0 ms) and 50 ms after stimulus onset, respectively. The definition of upper-L4 is indicated next to the CSD map.

We used current source density (CSD) analysis based on LFP signals (Mitzdorf, 1985) for identification of laminar boundaries, in particular the location of V1 input layer (L)4C, aided by spiking activity for identifying the top and bottom of the cortex. Specifically, CSD defined as the second spatial derivative of LFP signals, produces a map of current sinks (negative voltage fluctuations) and sources (positive fluctuations) as a function of time; this allows us to reliably identify, in response to stimulation of the neurons’ RFs, input L4C as the location of the earliest current sink followed by a reversal to current source (**Fig. 2C**) (Givre et al., 1995; Schroeder et al., 1998). The top of the cortex (L1) was identified as one contact (100μm) above the first contact where visually-driven spiking responses could be recorded, while the L6/white matter boundary was identified as the deepest contact at which visually-driven spike rates dropped by more than 50% compared to responses recorded in the contacts immediately above (**Fig. 2A**). The boundaries between other layers were estimated based on previous anatomical studies (Lund, 1973) and verified using postmortem histology, therefore, these boundaries are necessarily tentative. Specifically, we identified upper-L4 as a region extending from the uppermost part of L4C (i.e. upper layer 4Cα) to approximately 1-2 contacts above the top of 4C; this region likely encompassed layers 4B and upper-4Cα, and possibly layer 4A (**Figs. 1, 2C**). Layers 2/3 were identified as the layers between upper-L4 and L1, and the deep layers were the layers below L4C, within which we identified an upper (U) and lower (L) half as approximations for L5 and L6, respectively (**Figs. 1, 2B,C**). We also refer to all layers above L4C as superficial layers.

To investigate the laminar processing of surround signals, we recorded neuronal responses through a V1 column to visual stimuli presented at increasing distances from the RF of the column; this experimental design was motivated by previous studies implicating different circuits in the processing of surround stimuli located near versus far from the RF (Angelucci et al., 2002b, 2017). To identify the circuits carrying visual signals from the surround to the recorded V1 column, we recorded LFP responses to visual stimulation of the surround only, and calculated the onset latency of current sinks in the CSD across layers. In the absence of direct RF stimulation, surround stimuli do not evoke significant spiking responses from the recorded neurons, therefore the LFP largely reflects presynaptic activity and postsynaptic subthreshold responses evoked by surround stimulation (Buzsaki et al., 2012). This approach is better suited to localize laminar activation than measuring onset latency of spiking responses, because the dendritic arbors of cortical neurons, where most synaptic integration occurs, rarely co-localize with the soma, where spikes are initiated (Callaway, 2014). Instead, the best way to localize laminar activity with high spatial precision is to measure the extracellular current sinks that reflect the net post-synaptic potentials of local neurons (Givre et al., 1994; Schroeder et al., 1990, 1991; Tenke et al., 1993).

### Near and far surround stimuli evoke distinct laminar patterns of CSD signals

After mapping the location and size of the minimum response field (mRF) and the stimulus preferences of neurons across the recorded V1 column (see Methods), we recorded LFP and MUA in response to 0.5° black square stimuli flashed at increasing distances from the mRF of neurons in the column. Distances from the mRF center to the inner (i.e. closest to the mRF) edge of the square stimulus ranged between 0° and 1.25°, and the mRF surround was systematically probed. These small stimuli failed to evoke reliable LFP signals when presented beyond these distances. Therefore, we also used larger stimuli, consisting of static sinusoidal annular gratings (2° in width) flashed at distances from the mRF center ranging between 0.2° and 6° (measured from the mRF center to the inner edge of the annulus). This larger stimulus evoked robust LFP signals from the far surround, but we found it was too coarse to finely probe the near surround. To localize the LFP signals to specific layers, we performed CSD analysis and computed the onset latency of current sinks across layers (see Methods).

**Figure 3A-C** shows the half-waved rectified, baseline corrected (z-scored; see Methods) CSD (middle panels), and MUA (bottom panels) recorded in one example penetration in response to stimulation of the mRF, near surround and far surround (stimuli depicted in the top panels). We performed half-wave rectification of the z-scored CSD in order to eliminate current sources (which mostly reflect passive return currents) and focus on the time-varying sinks that indicate changes in excitatory synaptic activation. Small square stimuli inside the columnar mRF (**Fig. 3A** top panel) evoked the fastest current sink in L4C, followed by sinks in deep layers and then in superficial layers (**Fig. 3A** middle panel). The fastest evoked postsynaptic spiking activity, in response to this stimulus, occurred in L4C and mid-deep layers (**Fig. 3A** bottom panel). Early activation of L4C by RF stimulation can be explained by feedforward activation of this layer, where geniculocortical afferents predominantly terminate (**Fig. 1A**), and is consistent with previous findings (Schroeder et al., 1998).

**Figure 3.**
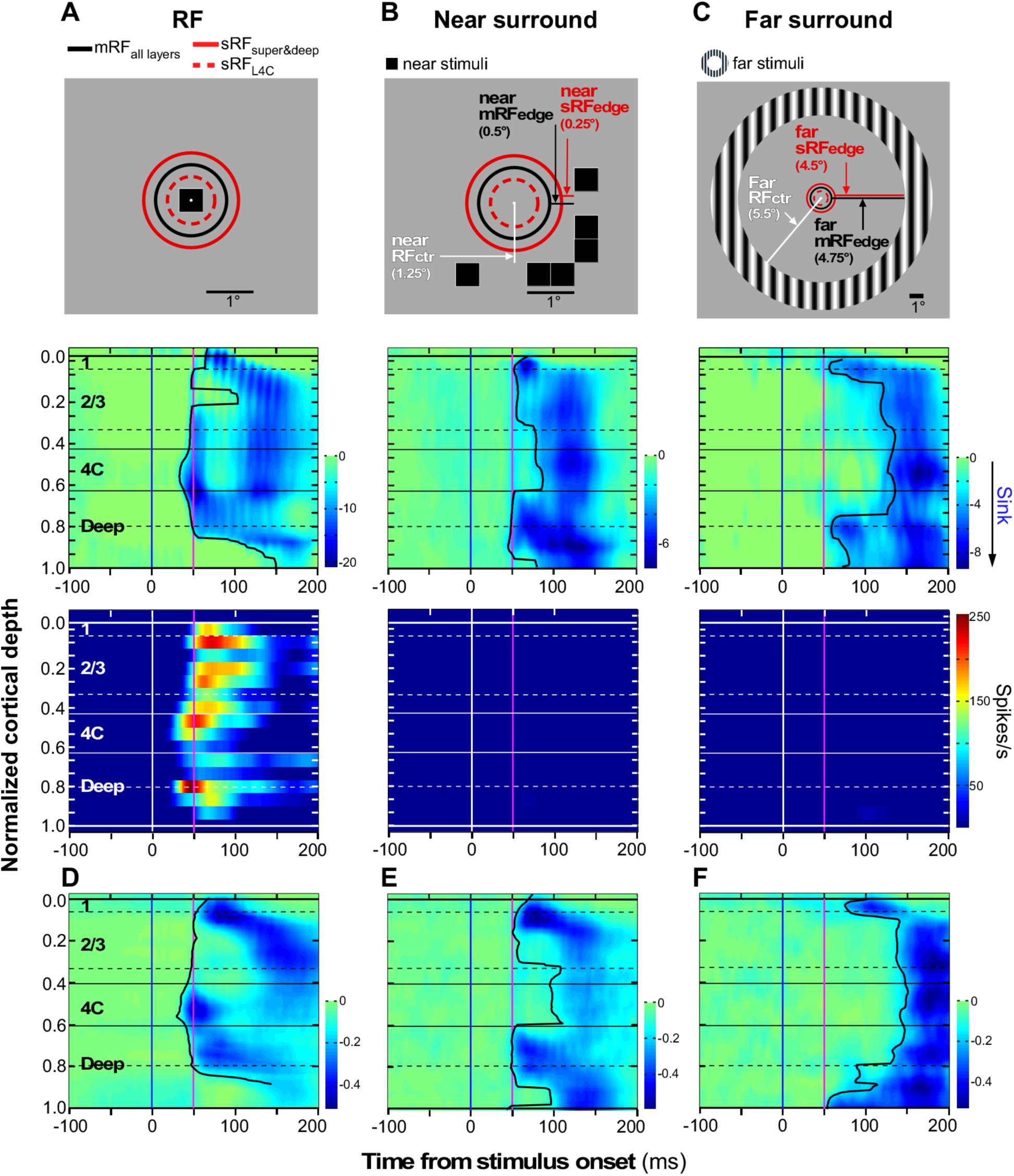
Laminar patterns of CSD and MUA signals evoked by stimulation of the RF, near or far surround. **(A-C) <u>Top panels</u>:** Location of the visual stimuli used (0.5° black squares or 2°-width annular gratings) relative to the mRF and sRF of the neurons recorded in one example penetration; the CSD and MUA responses to these stimuli in this penetration are shown in the middle and bottom panels, respectively. *Red solid circle:* largest sRF size measured across all layers; *red dashed circle:* largest sRF size measured in L4C; *black circle:* largest mRF size measured across all layers. **<u>Middle panels</u>:** Baseline-corrected (z-scored; see Methods) and half-wave rectified CSD signals (positive values are set to zero) recorded in the example penetration across V1 layers in response to presentation of the stimulus indicated in the panel at the top. The CSD profile in panel (B) is the average of 6 CSD profiles evoked by the small square stimulus presented at each of the indicated positions around the RF. Other conventions are as in **Fig. 2C**. **<u>Bottom panels</u>:** Color maps of MUA activity in response to the same stimuli for the same penetration. Color scale applies to panels (A-C). **(D-F)** Population averages of z-scored and half-wave rectified CSD evoked by stimulation of the RF (D), near surround (E) and far surround (F), using similar stimuli as shown in the top panels of (A-C). Near surround stimuli encompassed distances from the mRF center between 0.75° and 1.25°, and far surround stimuli between 1.42° and 5.5°.

As the stimulus was progressively moved away from the columnar mRF, spiking activity ceased first in L4C, then in superficial layers, and finally in deep layers, consistent with previous reports of larger mRFs in deep layers (Blasdel and Fitzpatrick, 1984; Gilbert, 1977; Hubel and Wiesel, 1977). When the stimulus reached a distance of just >1° from the mRF center (e.g. **Fig. 3B** top panel), CSD signals in L4C were delayed, and the earliest current sinks were observed in the superficial and deep layers almost at the same time (the lower half of the deep layers was activated first in the specific example case of **Fig. 3B**). To understand how the location of the surround stimulus related to the boundaries of the RF of neurons in the recorded column, and because RF size varies depending on the methods used to map it (Angelucci and Bressloff, 2006; Angelucci et al., 2002b), we measured the size of the mRF as well as of the summation RF (sRF: see Methods for mapping and definitions) across contacts. In the example recording of **Fig. 3B**, the inner edge of the surround stimulus was located 1.25° from the center of the columnar RF, corresponding to 0.5° outside the edge of the largest mRF recorded across the column (black circle in **Fig. 3B** top panel), or 0.25° outside the edge of the largest sRF in the column (solid red circle in **Fig. 3B** top panel). This near surround stimulus did not evoke significant spiking responses across the column (**Fig. 3B** bottom panel), therefore the CSD sinks it evoked reflected subthreshold responses. This laminar pattern of CSD signals evoked by small stimuli in the near surround suggests involvement of multiple connection types in the processing of local context, including horizontal connections in superficial and deep layers, and possibly feedback circuits (see Discussion).

Stimulation of the far surround with an annular grating located at distances >1.4° from the mRF center evoked the earliest CSD signals in feedback-recipient layers, i.e. L1 and the lower half of the deep layers, with much increased latency of signals in the remainder of the layers (**Fig. 3C** middle panel). In **Fig. 3C**, the inner edge of the surround annulus was located 5.5° from the mRF center, corresponding to 4.75° and 4.5° outside the outer edge of the largest mRF and sRF in the column, respectively. This laminar pattern of CSD signals suggests that visual signals in the far surround are relayed to the recorded V1 column by feedback connections (see Discussion).

The CSD grand averages for all recorded penetrations (**Fig. 3D-F;** see Methods) resembled the profiles of the example cases shown.

**Figure 4** shows the quantitative analysis of onset latency of current sinks across layers for all penetrations (n=10), including total of 39 stimulus conditions. To determine which layers were first activated by each surround stimulus condition, within a penetration for each stimulus condition we computed a relative latency (ΔL) as the difference between the onset latency of currents sinks at each contact and the shortest latency across all contacts. ΔL values were, then, pooled across all conditions and penetrations by layer location of the contact, and population means and medians were computed (**Fig. 4A-C** left column). Lower values of ΔL for a layer indicate shorter onset latency in that layer.

**Figure 4.**
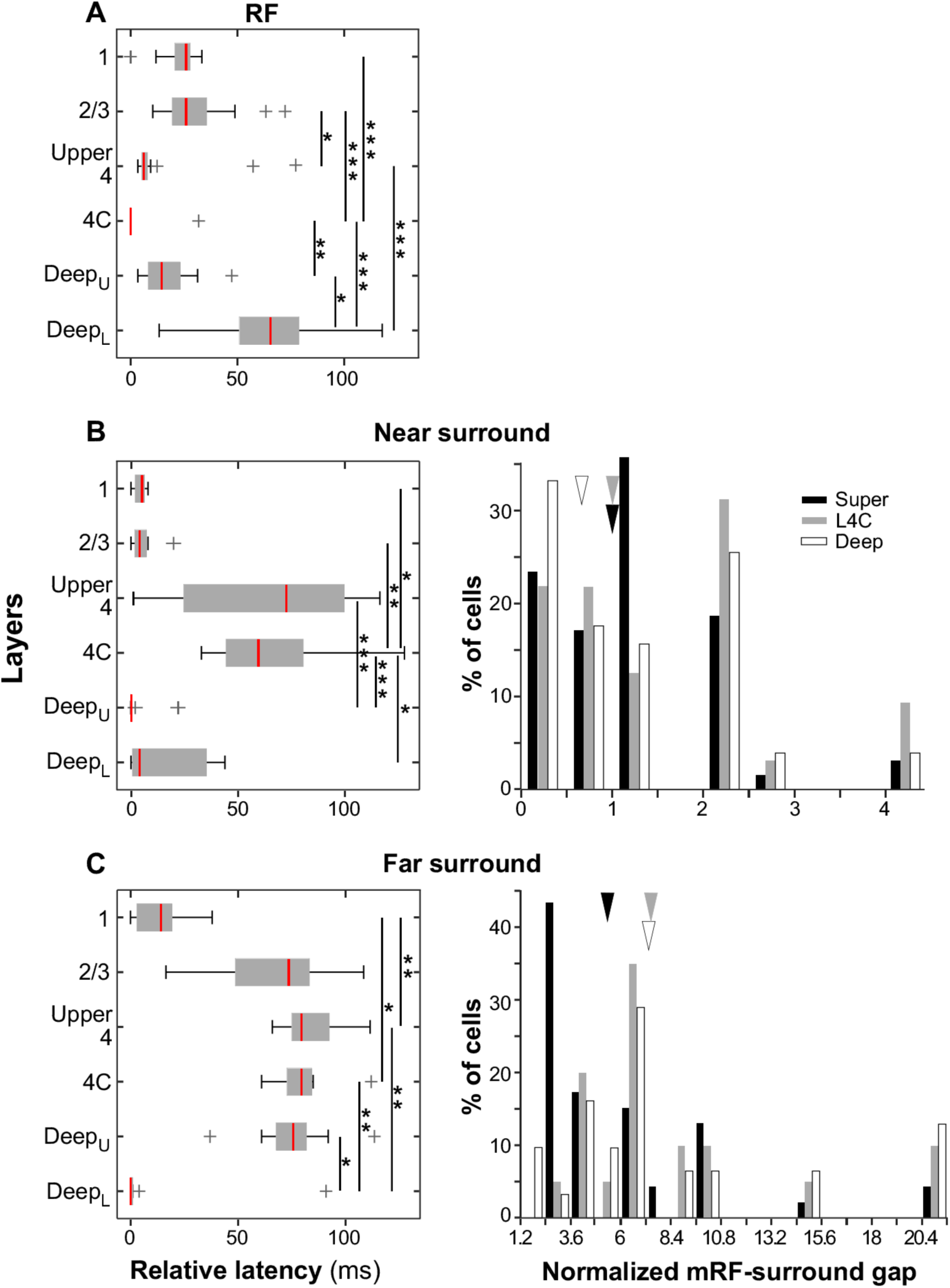
Onset latency of CSD signals across layers evoked by stimulation of the RF, near or far surround. **(A-C) <u>Left panels</u>:** Box plots of relative latency in response to stimuli in the RF (A), near (B) or far (C) surround. *Red vertical lines:* median values. (**B-C) <u>Right panels</u>**: Distributions of normalized mRF-surround gaps for stimuli in the near surround (B; n= 147 contacts) and far surround (C; n= 97 contacts). Different color bars indicate different layers. *Arrowheads:* median latency.

Stimuli inside the RF evoked the earliest current sinks in L4C (mean ΔL±s.e.m, 1.88±1.88 ms, n=17 conditions); onset latency in this layer was significantly shorter than in L1 (23.32±2.06 ms, n=17, p<0.0001; Kruskal-Wallis test with Bonferroni correction), L2/3 (29.85±4.18 ms, n=17, p<0.0001) and the upper and lower halves of the deep layers (Deep_U_: 17.23±2.63 ms, n=17, p=0.0053; Deep_L_: 62.57±7.31 ms, n=17, p=<0.0001), but it was not significantly different from latency in upper-L4 (13.55±5.02 ms, n=17. p=0.43) (**Fig. 4A**).

In contrast, stimuli in the near surround (i.e. located at distances from the mRF center between 0.75° and 1.25°) evoked delayed current sinks in L4C (66.72±8.39 ms, n=13), and onset latency in this layer was significantly longer than in all other layers (L1: 4.5±0.74 ms, n=13, p=0.0174; L2/3: 4.92±1.46 ms, n=13, p= 0.0069; Deep_U_: 2.18±1.83 ms, n=13, p<0.0001; Deep_L_: 16.5±5.19 ms, n=13, p=0.043), except upper-L4 (63.11±8.4 ms, n=13, p=1.0) (**Fig.4B** left panel). Despite a tendency for the upper half of the deep layers to have the shortest latency, onset latency in this layer was only significantly earlier than in L4C (p<0.0001) and upper-L4 (p<0.001), but not the remaining layers (p>0.05). Upper-L4 also did not differ in latency from other layers, except Deep_U_. The right panel of **Fig. 4B** shows the location of the near surround stimuli relative to the edge of the mRF of neurons in different layers. This is expressed as “normalized mRF-surround gap”, defined as

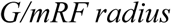

where *G* is the gap (in degrees) between the inner edge of the surround stimulus and the edge of the mRF, and *mRF radius* is the radius of the mRF. Thus, a normalized mRF-surround gap equal to zero indicates that the inner edge of the surround stimulus abutted the mRF edge (there was no gap), negative values indicate that the surround stimulus was inside the mRF, and positive values that the stimulus was located outside the mRF; for example, a value of 2 indicates that the gap between the mRF edge and the surround stimulus was twice the mRF radius. The distribution of normalized mRF-surround gaps in **Fig. 4B** ranges from 0 to 4, with median values of 1 (for superficial layers and L4C) and 0.67 (for deep layers); this indicates that the stimuli were located outside the mRF of neurons across the whole column. In a similar fashion, we measured the location of the near surround stimuli relative to the edge of the sRF of the same neurons, by estimating a “normalized sRF-surround gap”; median gap values were 0.67 for superficial layers (range= -0.1 to 4) and for deep layers (range= -0.3 to 3), and 1 for L4C (range= 0 to 3) (see **Supplementary Fig. 1A**). Thus for most contacts the near surround stimuli were also located outside the sRF of L4C neurons in the column, but for a few contacts in the superficial and deep layers (11/138) they activated the fringes of the sRF. To ascertain that the CSD profile observed in response to near surround stimuli did not reflect direct stimulation of neuronal sRFs by the stimuli intended to stimulate the surround, we performed the same onset latency analysis after discarding stimulus conditions with negative values of normalized sRF-surround gap, and found similar CSD profiles and quantitative results as in **Figs. 3E and 4B (**see **Supplementary Fig. 1B-C**).

Stimuli in the far surround (i.e. located at distances from the mRF center >1.4°) evoked the earliest current sinks in L1 (mean ΔL±s.e.m, 13.72±4.22 ms, n=9), and the lower half of the deep layers (10.56±10.06 ms, n=9); onset latencies in L1 and Deep_L_ were significantly shorter than in L4C (80.5±4.99 ms, p= 0.0186 and 0.0049, respectively) and upper-L4 (83.93±4.8 ms, p= 0.007 and 0.0017, respectively), and latency in Deep_L_ was significantly shorter than in Deep_U_ (75.33±6.93 ms, p=0.0227) (**Fig. 4C** left panel). The normalized mRF-surround gap for these stimuli had median values of 4.27 (superficial layers, range 2.4-21), 6.4 (L4C, range 2.6-21) and 6.3 (deep layers, 1.2-21) (**Fig. 4C** right panel); the normalized sRF-surround gap had median values of 6.6 (superficial layers, range 3.4-27), 7 (L4C, range 2.16-12.75), and 7 (deep layers, range 0.9-12.75; see **Supplementary Fig. 2**), indicating all far surround stimuli were located well outside both the mRF and sRF of the recorded neurons.

In summary, stimuli in the near and far surround of the recorded V1 column evoked different laminar patterns of CSD signals, suggesting involvement of different circuits and layers, in the processing of local and global context, respectively.

### Onset latency of near and far surround suppression across V1 layers

The results presented above point to the circuits that carry near and far surround signals to the V1 center column; surround suppression is likely to be initiated in the layers where these circuits terminate and evoke the earliest postsynaptic responses. To understand how these surround pathways contribute to surround suppression in different layers, we presented visual stimuli simultaneously in the RF and surround, and measured the onset latency of surround suppression of spiking responses (MUA) across layers (see Methods). We used oriented grating stimuli, for these experiments, because we wanted to determine the onset latency of both orientation tuned and untuned suppression; moreover, the small square stimuli did not evoke sufficiently strong suppression to allow for reliable latency measurements, and often evoked facilitation [consistent with previous studies (see Angelucci and Shushruth, 2013 for a review)]. We used two kinds of grating stimuli (see Methods): a 20° diameter grating patch centered on the columnar RF and presented at the optimal orientation for the recorded V1 column (**Fig. 5A**, top inset); and 2°-width annular gratings of optimal or orthogonal-to-optimal orientations flashed at increasing distances from the RF (as described above) and presented together with grating patches of optimal orientation and size for the columnar RF (**Fig. 5B-C** top insets).

**Figure 5.**
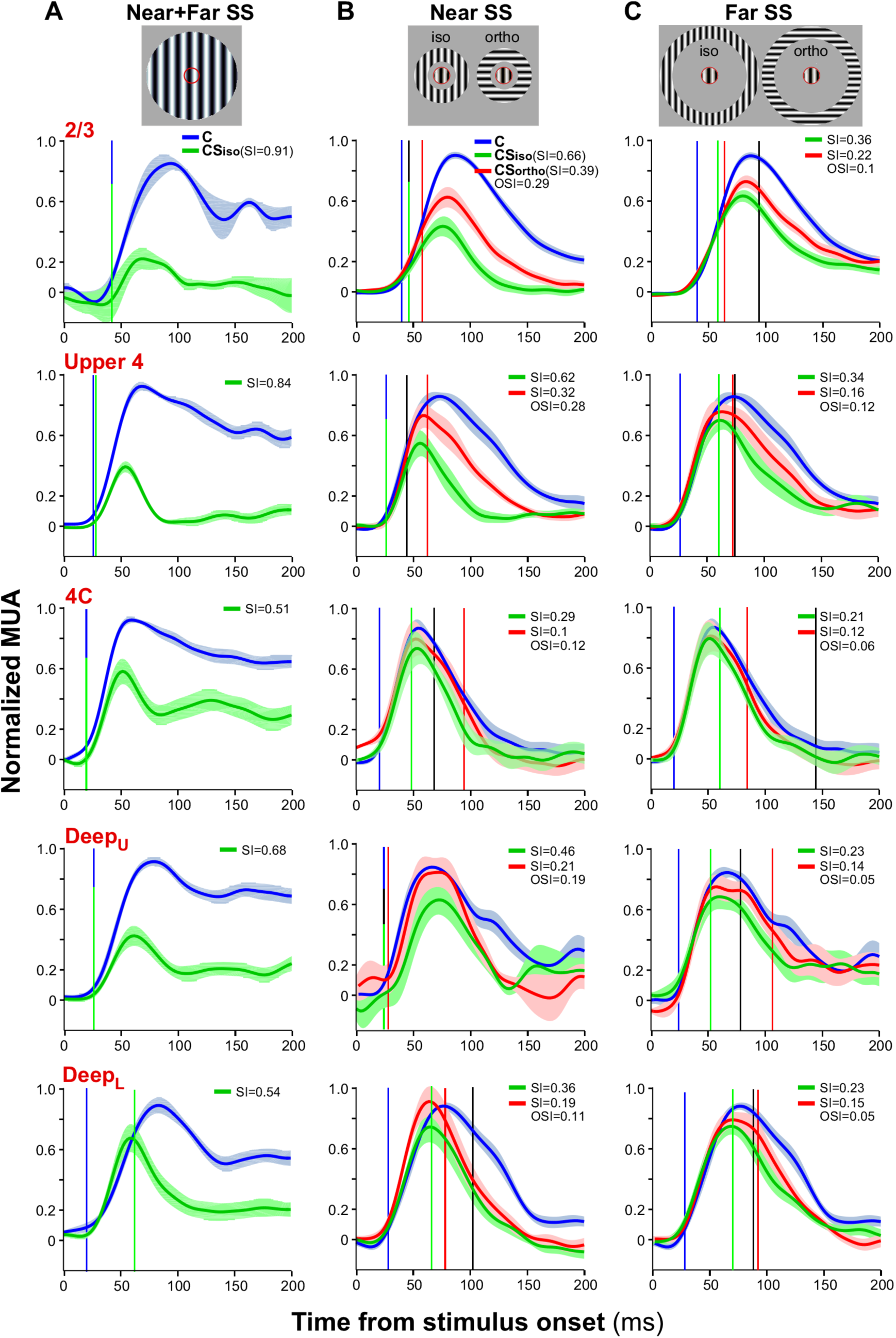
Laminar patterns of MUA responses evoked by center-only and center-surround gratings. **(A-C) <u>Top row</u>:** Center-surround grating stimuli used to probe near and far surround suppression across the recorded V1 column. *Red circle* outlines the size of the aggregate sRF of neurons in the column. **<u>Bottom five rows</u>:** averaged PSTHs in response to the center-only stimulus (*blue*) and the respective center-surround stimuli shown at the top of each column. *Green curves:* responses to iso-oriented center-surround gratings; *red curves:* responses to orthogonally-oriented center-surround gratings. The response at each contact was normalized to the peak of the center-only response in the first 300ms, and then responses within each layer were averaged. *Shade:* ±1 sem. *Vertical lines* indicate the onset latency of responses (*blue:* center-only; *green:* iso-oriented suppression; *red:* orthogonally-oriented suppression; *black:* tuned suppression). SI values indicate the suppression index caused by iso-oriented and ortho-oriented surround stimuli, and the OSI the orientation-selectivity index of the suppression, measured from the averaged PSTHs.

**Figure 5** shows the peristimulus time histograms (PSTHs) for the population of penetrations, averaged by layer, in response to the center-only stimulus (*blue curve*), iso-oriented or orthogonally-oriented center-surround stimuli (*green and red curves*, respectively). In each panel, we report the strength of surround suppression, expressed as a suppression index (SI; see Methods) estimated from the average responses, and ranging from 0 (no suppression) to 1 (complete suppression); the panels of **Fig. 5B-C** also report the orientation selectivity of surround suppression, expressed as an orientation selectivity index (OSI; see Methods) measured from the average responses, and ranging from 0 (no selectivity) to 1 (high selectivity). The data in **Figure 5** confirm several previous findings on the strength and orientation tuning of surround suppression: 1. in all layers SI caused by iso-oriented surround stimuli decreases with increasing distance of the surround stimulus from the RF (Sceniak et al., 2001; Shushruth et al., 2009); 2. SI is greater above L4C and smaller in L4C and deep layers (Ichida et al., 2007; Sceniak et al., 2001; Shushruth et al., 2009); 3. in all layers, SI is generally greater for iso-oriented than for orthogonally-oriented center-surround stimuli, i.e. surround suppression is orientation-tuned, and this is true for both near and far surround suppression, although far surround suppression in all layers is less tuned than near suppression (Shushruth et al., 2013); 4. Both near and far surround suppression are more sharply tuned (larger OSI values) above L4C (Henry et al., 2013; Shushruth et al., 2013).

We measured the onset latency of surround suppression as the time point at which the MUA profile in response to the center-surround stimuli diverged significantly from the MUA profile in response to the center-only stimulus (see Methods). In **Figure 5** onset latency was measured on the layer-averaged responses for our entire cell sample. In response to the large (20° diameter) grating patch, the earliest suppression occurred in L4C and at the same time as the onset latency of the center-only response (**Fig. 5A**); in other layers, although the latency of suppression was longer, it also coincided with the onset latency of the center-only response. These findings strongly suggest that the earliest suppression in V1 is relayed from LGN. An exception was the lower half of the deep layers, in which the onset of suppression occurred on average 42 ms after onset of the center response, but early suppression in this layer may have been masked by an early facilitation seen in the average response profile.

When an iso-oriented surround stimulus was moved away (>1.1°) from the RF center into the near surround (**Fig. 5B**), the onset of suppression in L4C was markedly delayed relative to the suppression onset caused by the large patch stimulus, and also relative to other layers (except Deep_L_). This observation, and the fact that near-surround suppression was weakest in L4C indicate that the suppression caused by stimuli in the near surround is generated intracortically, outside L4C. In response to stimuli in the near surround, the earliest suppression occurred in upper-L4, where horizontal connections within the V1 column first occur as visual signals are relayed outside L4C. Fast suppression also occurred in the upper half of the deep layers, although a cell-by-cell analysis demonstrated variability in the onset latency of suppression in this layer (see below).

We also measured the onset latency of tuned surround suppression as the time point at which the response curves to iso-oriented and cross-oriented center-surround stimuli diverged significantly (*black line* in **Fig. 5B;** see Methods). It is noteworthy that tuned near surround suppression in L2/3 and Deep_U_ appeared on average at the same time as the earliest suppression, suggesting an involvement of orientation-specific horizontal connections in these layers. In contrast, tuned suppression in upper-L4 was delayed relative to the earliest suppression, suggesting tuned suppression in this layer may be relayed from other layers.

The averaged population responses in **Figure 5C** indicate that far surround stimuli (>2° from RF center) evoked the earliest suppression in the upper part of the deep layers, but since many contacts showed weak-to-absent far surround suppression in L4C and deep layers, we performed a contact-by-contact analysis of onset latencies based only on cells that showed significant suppression (see below).

**Figure 6** shows the quantitative analysis of onset latency of surround suppression across layers for our population of penetrations. For each contact, we estimated the onset latency of surround suppression as the time point at which the MUA profile in response to the center-surround stimulus diverged significantly from the MUA profile in response to the center-only stimulus; the onset latency of tuned suppression, instead, was defined as the time point at which the MUA profiles in response to iso- and ortho-center-surround stimuli diverged from each other. We included in this analysis only cells that showed significant suppression (SI>0.15) (see Methods). To determine in which layers suppression emerged earliest in time, for each contact within a penetration we computed a relative latency (ΔL) as the difference between onset latency at that contact and the shortest latency across all contacts in the same penetration for any given center-surround stimulus. ΔL values were, then, pooled across penetrations by layer location of the contact, and population means and medians were computed (**Fig. 6**).

**Figure 6.**
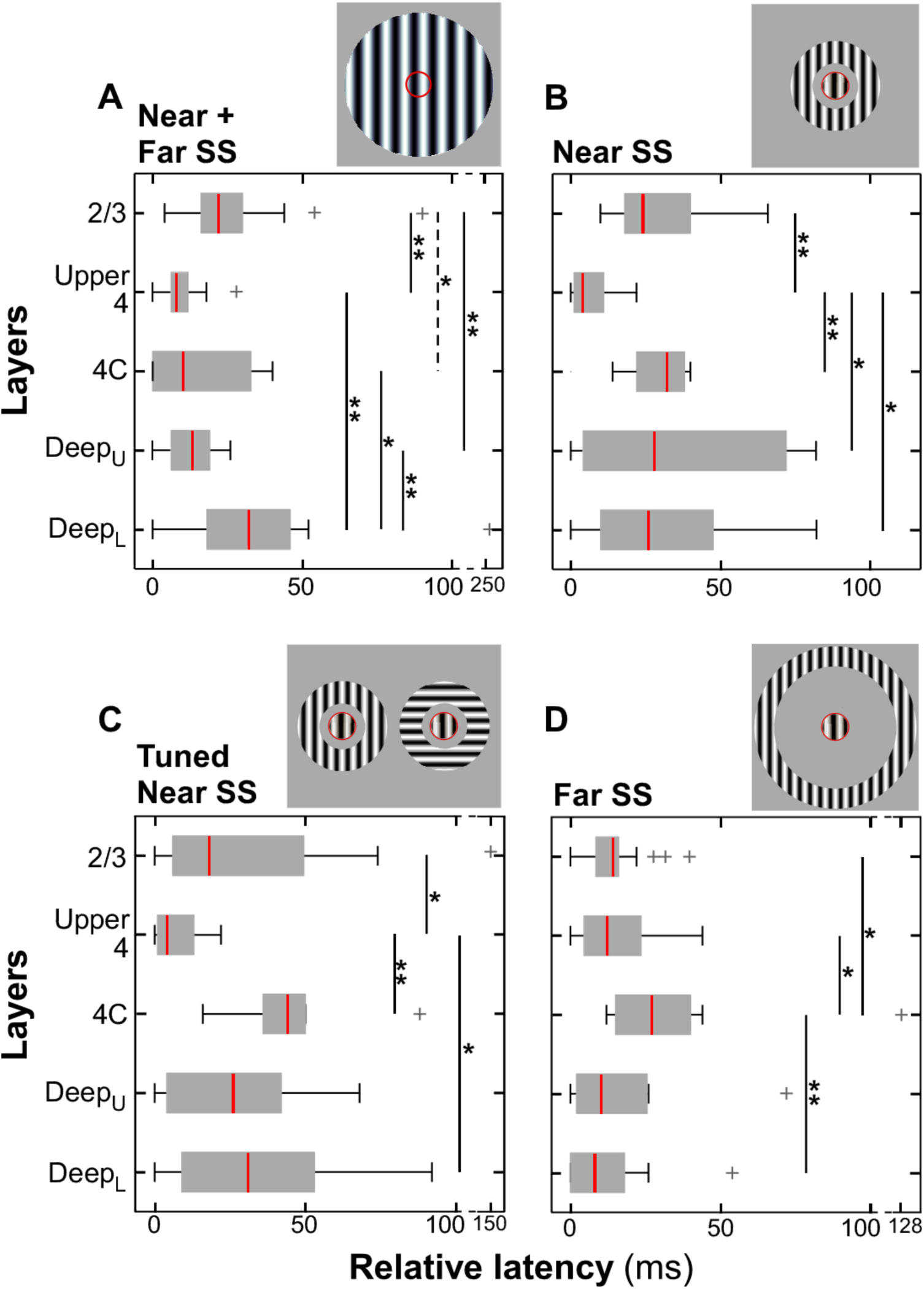
Onset latency of surround suppression across layers evoked by center-surround gratings. **(A-D)** Box plots of relative surround suppression latency measured in response to a 20° diameter grating patch encompassing the RF, near and far surround (A; n=6 penetrations), a center grating patch and annular grating encompassing the RF and near surround (B-C; n=4 penetrations), or a center grating patch and annular grating encompassing the RF and far surround (D; n=6 penetrations). (A,B,D): onset latencies in response to iso-oriented center-surround stimuli; (C) onset latency of tuned suppression. *Red vertical lines:* median values. Dashed vertical line in (A) indicates p=0.05.

Large grating patch stimuli encompassing the RF, near and far surround, evoked the earliest suppression in L4C (mean ΔL±s.e.m: 16.18±4.15 ms, n=15 contacts), upper-L4 (10.01±2.43 ms, n=11) and Deep_U_ (12.17±2.21 ms, n=14), with no significant difference in the latency of suppression between these layers (**Fig. 6A**). Median latency was fastest in L4C (10.01 ms) and upper-L4 (8 ms), but not significantly faster than in Deep_U_ (13.01 ms), due to a larger scatter of onset latencies in L4C. Onset latencies in L4C, upper-L4 and Deep_U_ were significantly different from latency in Deep_L_ (median: 32.03, mean±s.e.m.: 45.12±14.81 ms, n=19; p=0.023, 0.001, 0.003, respectively; Kruskal-Wallis test with least significant test correction) and in L2/3 (median: 22.02 ms, mean±s.e.m.: 25.61±3.02 ms, n=29; p=0.055, 0.002, 0.006, respectively).

In contrast, iso-oriented stimuli presented in the near surround, but at distances>1° from the RF center, evoked the earliest suppression in upper-L4 (median: 4 ms,
mean±s.e.m.:6.75±2.72 ms, n=8), followed by L2/3 (median: 24.02 ms, mean±s.e.m.: 29.5±3.57 ms, n= 21), and suppression in upper-L4 was significantly faster than in all other layers (L2/3: p=0.0018; Deep_L_: median: 26.02 ms, mean±s.e.m.: 30.69±8.78 ms, n=10, p=0.016; Deep_U_: median: 28.02 ms, mean±s.e.m.: 35.7±14.58 ms, n=6, p=0.029; L4C: median: 32.03 ms, mean±s.e.m.: 29.69±3.41 ms, n=9, p=0.0082) (**Fig. 6B**). L4C showed delayed suppression, whereas suppression latency showed variability in the deep layers with cells in these layers showing both fast as well as delayed suppression onset. The onset latency of tuned near surround suppression showed a similar laminar profile as the latency of iso-oriented near surround suppression, emerging first in upper-L4 (median: 4 ms, mean±s.e.m.: 7.25±2.95 ms, n=8), followed by L2/3 (median: 18.01 ms, mean±s.e.m.: 32.13±7.87 ms, n=21), with latency in upper-L4 being significantly faster than in all other layers (L2/3: p=0.025; L4C: median: 44.04 mean±s.e.m.: 46.37±7.92 ms, n=9, p=0.004; Deep_L_: median: 31.02, mean±s.e.m.: 34.78±10.06 ms, n=10, p=0.048), except Deep_U_ (median: 26.02 ms, mean±s.e.m: 27.69±10.39 ms, n=6, p=0.11) (**Fig. 6C**).

Iso-oriented far surround stimuli evoked the earliest suppression almost simultaneously in all layers, except L4C (median: 27.02 ms, mean±s.e.m.: 38.03±10.17 ms, n=14 contacts) in which onset latency was significantly delayed compared to upper-L4 (median: 12.01 ms, mean±s.e.m.: 15.28±4.13 ms, n=11, p=0.042), L2/3 (median: 14.01 ms, mean±s.e.m.: 14.16±1.69 ms, n=32, p=0.021) and Deep_L_ (median: 8 ms, mean±s.e.m.: 12.29±3.62 ms, n=17, p=0.005), but not compared to Deep_U_ (median: 10 ms, mean±s.e.m.: 19.44±7.69 ms, n=11, p =0.053) (**Fig. 6D**). Notably, many contacts in L4C did not show significant far surround suppression, therefore these cells did not contribute to the average onset latency in this layer; lack of far suppression in L4C (see also **Fig. 5C**) further supports the notion that far suppression is generated outside this layer, which lacks feedback (and horizontal) connections. We did not measure the onset latency of tuned far surround suppression as significant tuning was observed only in L2/3 and upper-L4. In summary, near and far surround suppression showed different laminar profiles of onset latencies. There was no consistent difference in absolute latencies between near and far surround suppression, but the two were positively and significantly correlated (see **Supplementary Fig. 3**).

## III. DISCUSSION

We have examined how the different cortical layers integrate visual information arising from outside neuronal RFs and contribute to the emergence of surround suppression. In macaque V1, we found layer-specific differences in the onset latency of subthreshold responses and of suppression of visually-evoked spiking responses induced by stimuli in the near versus far surround. These laminar differences corresponded well with the anatomy of feedforward, horizontal and feedback connections (**Fig. 1**).

### Laminar Processing of Stimuli in the Receptive Field

Stimuli inside the RF evoked the earliest current sinks in L4C, consistent with initial activation of this layer by feedforward afferents from the LGN, which terminate predominantly in L4C (Lund, 1988). However, the earliest spiking activity following RF stimulation occurred in both L4C and deep layers. It is unclear why early spiking activity in the deep layers was not reflected in early current sinks in these layers. One possibility is that the sparse LGN inputs to L6 in macaque V1 (Lund, 1988) are too weak to drive L6 neurons. However, deep layer neurons could be activated by LGN inputs into L4C, where the apical dendrites of these neurons reside (Lund, 1988). This would result in early subthreshold responses in L4C followed by spiking activity in both L4C and deep layers, which is what we observed.

### Laminar Processing of Stimuli in the Near Surround

Stimulation of the surround region just outside the columnar RF, i.e. the near surround, without simultaneous stimulation of the RF, evoked the earliest current sinks in the superficial and deep layers almost at the same time. Current sinks in L4C were delayed, thus ruling out any involvement of geniculocortical afferents to L4C in the processing of near surround signals, and suggesting these signals are conveyed to the center column via intracortical connections outside L4C. Early activation of all layers, but 4C, suggests near surround stimuli recruit multiple circuits, including intra-V1 horizontal connections and possibly inter-areal feedback connections. Horizontal connections are present in all layers, except L1 and mid-to-lower L4C, while feedback connections are strongest in L1 and the lower half of the deep layers, and absent in L4C (**Fig. 1**). Therefore, early subthreshold activation of L1 by near surround stimuli implicates involvement of circuits other than horizontal connections, such as feedback, as also suggested by early and strong activation of the lower deep layers where feedback terminations are denser than horizontal connections (Federer et al., 2015; Lund et al., 2003; Rockland and Pandya, 1979). This would suggest that feedback connections are extremely fast and can conduct signals to the center column almost as fast as intra-V1 horizontal connections. Indeed, we also found similar onset latencies for near and far surround suppression (**Supplementary Fig. 3**). This conclusion is consistent with previous studies demonstrating that orientation-tuned surround suppression can be delayed by as little as 10 ms relative to the onset of visually evoked responses in V1, and that onset latency of suppression is independent of the distance of the surround stimulus from the RF (Bair et al., 2003). However, it is unclear whether the small 0.5° square stimuli are sufficiently large to significantly activate extrastriate cortex, and therefore feedback neurons.

Alternatively, or in addition, early current sinks in L1 following near surround stimulation could result from recruitment of geniculocortical K1-K2 koniocellular afferents. The latter have terminations in L1 that are spatially more widespread (up to 1 mm) (Casagrande et al., 2007; Lund, 1973) than those of geniculate magno- or parvo-cellular afferents in L4C and 6 (Angelucci and Sainsbury, 2006; Lund, 1973, 1988), or those of K3-K6 koniocellular afferents in L2/3 (Casagrande et al., 2007).

Surround suppression of spiking responses induced by large stimuli encompassing the RF and the full extent of the surround first emerged in L4C, and at the same time as the onset of responses to stimuli confined to the RF. This strongly suggests that the earliest suppression in V1 is inherited from the LGN; namely, the large stimuli cause surround suppression of LGN cells (Alitto and Usrey, 2008; Sceniak et al., 2006), resulting in withdrawal of feedforward excitation to V1 cells.

Introducing a small gap between the stimulus in the RF and that in the near surround, so that the surround stimulus was likely located beyond the anatomical spread of geniculocortical afferents to the center column, led to delayed suppression in L4C relative to other layers, pointing to an intracortical origin of the suppression, outside L4C. Under this stimulus condition, both the earliest suppression and orientation-tuned suppression first emerged in the superficial layers, particularly in upper-L4 (encompassing L4A to upper-4Cα). L4B and upper-4Cα share a system of horizontal connections that is the first long range system to appear along the flow of visual information exiting L4C (Angelucci et al., 2002a; Lund et al., 2003). Therefore, it is possible that horizontal connections in upper-L4 are activated by near surround stimuli earlier than horizontal connections in downstream layers. It is, however, curious that we did not observe consistent early current sinks in upper-L4, but rather a wide distribution of onset latencies, when stimuli were presented to the near surround only. Perhaps, because horizontal (and feedback) connections in upper-L4 are weaker than in other layers, it is more difficult to activate them, or their activation induces weak current sinks, which our latency analysis could not capture. Alternatively, since the apical dendrites of pyramids in L4B reach the superficial layers, surround suppression of these cells could be generated through their apical dendrites, by disynaptic inhibition initiated by horizontal and/or feedback connections within superficial layers.

That orientation-tuned suppression emerges first in the superficial layers is consistent with the orientation-specific organization of horizontal connections previously described for these layers (Malach et al., 1993) (albeit an orientation-specific organization remains to be demonstrated for horizontal connections in upper-L4), and suggests that the deeper layers may, instead, inherit tuned suppression from the superficial layers.

### Laminar Processing of Stimuli in the Far Surround

Stimulation of the far surround without simultaneous stimulation of the RF evoked the earliest current sinks in L1 and the lower half of the deep layers, where feedback terminations are particularly dense. This suggests that visual signals in the far surround are relayed to these layers in the center column via feedback connections, and therefore that far surround suppression is initiated by feedback. Far surround suppression emerged first and almost simultaneously in superficial and deep layers, and last in L4C, a layer that lacks both horizontal and feedback connections, and within which neurons largely confine their dendrites. Since stimuli in the far surround evoked the earliest current sinks in L1/2A and lower deep layers, early suppression in the superficial and deep layers must be initiated by feedback contacts with inhibitory cells in L1/2A and 5/6. In turn, inhibitory neurons in L1-2A can suppress pyramidal cells in most layers via contacts with these cells’ apical dendrites ascending to L1. Pyramids in L2-4B have apical dendrites that often ascend to L1, while pyramids in L5-6 do not consistently send apical dendrites to L1, and L4C lacks pyramidal cells and its neurons confine their dendrites largely to 4C (Callaway and Wiser, 1996; Lund, 1973). Inhibitory neurons in L5/6, thus, could suppress neurons in these layers that do not send apical dendrites to L1/2A, while L4C could inherit late far surround suppression from other layers. The synaptic mechanisms that may generate surround suppression are reviewed in detail in Angelucci et al. (2017)

### Relationship to Prior Studies

Our results indicate that near surround suppression results from the interaction of feedforward, horizontal and feedback connections, while far surround suppression is initiated by feedback connections. These results are consistent with a plethora of previous anatomical, physiological and optogenetic studies, and with our previously formulated hypothesis on the circuits for surround suppression and contextual interactions (these studies and hypothesis are reviewed in Angelucci et al., 2017; Angelucci and Bressloff, 2006; Angelucci and Shushruth, 2013).

The early studies of Angelucci et al. (2002b) demonstrated that monosynaptic horizontal connections are spatially co-extensive with the near surround of V1 neurons, while feedback connections encompass the full extent of the RF and surround, and led to the hypothesis illustrated in **Fig. 7**. Later studies demonstrated that LGN neurons show untuned surround suppression (Alitto and Usrey, 2008; Sceniak et al., 2006), and that untuned surround suppression in V1 can occur as fast as visual responses to RF stimulation (Henry et al., 2013), implicating LGN afferents in untuned suppression in V1. These findings are consistent with our results of earliest suppression in L4C.

**Figure 7.**
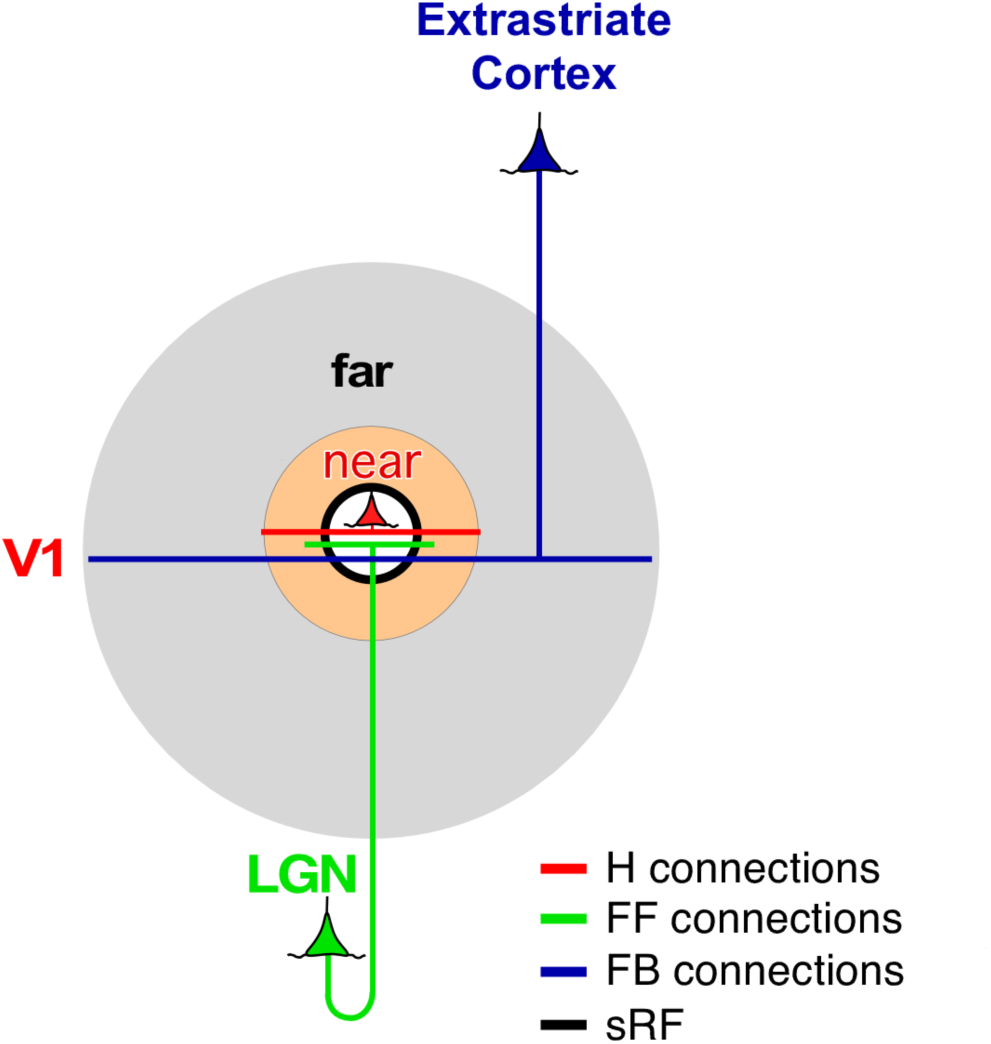
Circuits for surround suppression. Feedforward (FF; *green*), horizontal (H; *red*), and feedback (FB; *blue*) connections all contribute to the sRF (*white shaded area*) and to the near surround (*orange shaded area*), but only feedback connections contribute to the far surround (*gray shaded area*).

Other studies found that tuned suppression is delayed by 10-30 ms relative to the onset of visually-evoked RF responses and of untuned suppression, and is independent of the distance of the surround stimulus from the RF (Bair et al., 2003; Henry et al., 2013). These studies implicated intracortical circuits and feedback connections in generating tuned surround suppression. Additional studies demonstrated that the conduction velocity of horizontal axons, at least in superficial layers, is too slow to mediate far surround suppression, and suggested the latter is mediated by much faster conducting feedback axons (Girard et al., 2001).

Two recent optogenetic studies in mouse V1 have provided direct evidence for a role of horizontal connections in surround suppression (Adesnik et al., 2012; Sato et al., 2014). Instead, studies in which feedback activity was abolished by cooling or pharmacologically blocking an entire extrastriate area have provided contrasting results regarding the role of feedback in surround suppression. Some studies observed weak reduction of surround suppression after cooling primate area MT (Hupé et al., 1998) or V2 and V3 together (Nassi et al., 2013), or cat postero-medial temporal visual cortex (Bardy et al., 2009); others found general reduction in response gain but no changes in surround suppression after pharmacologically silencing primate V2 (Hupé et al., 2001), cooling cat postero-temporal visual cortex (Wang et al., 2010) or optogenetically silencing mouse cingulate cortex (Zhang et al., 2014). A recent study, in which the activity of feedback neurons from V2 was selectively reduced with varying intensity, found that moderate reduction of feedback activity reduced surround suppression and increased RF size, while strong reduction of feedback activity additionally caused a reduction in overall response gain in V1 (Nurminen et al., 2017).

## Author Contributions

M.B., L.N. and A.A. designed project. All authors collected electrophysiological data. M.B. analyzed electrophysiological data. S.M. analyzed histological sections for array penetration and generated histological figure. A.A. supervised all aspects of project. M.B. and A.A. wrote the paper, L.N. and S.M. edited the manuscript. All authors discussed the results, commented on and approved the final manuscript.

## Acknowledgments

We thank Kesi Sainsbury for technical assistance, and Drs. Frederick Federer and Jeff Yarch for help with experiments. This work was supported by grants from the National Institute of Health (R01 EY026812, R01 EY019743, BRAIN U01 NS099702), the National Science Foundation (IOS 1355075, EAGER 1649923), the University of Utah Research Foundation, The University of Utah Neuroscience Initiative, to A.A., a grant from Research to Prevent Blindness, Inc. to the Department of Ophthalmology, University of Utah, and a postdoctoral fellowship from the Ella and Georg Ehrnrooth Foundation to L.N.

## METHODS

### Experimental Model and Subject Details

Macaque monkeys (*Macaca fascicularis*) were purchased from a commercial breeder, quarantined for 6 weeks and group-housed at the University of Utah prior to being used for experimentation. Linear array recordings (total of 22 penetrations from 7 animals) were made in the parafoveal representation of the primary visual cortex (V1) in adult macaque monkeys (3-4 kg). We selected for analysis only those penetrations (n=10 from 4 macaques, 2 males and 2 females) that were deemed to be perpendicular to the surface of V1 according to the criteria described below. All experimental procedures were in accordance with protocols approved by the University of Utah Institutional Animal Care and Use Committee and with NIH guidelines.

### Methods Details

#### Surgical Procedures

Animals were pre-anesthetized with ketamine (25 mg/kg, i.m.), intubated, artificially ventilated with a 70:30 mixture of N_2_O and O_2_, and their head was fixed by positioning in a stereotaxic apparatus. During surgery, anesthesia was maintained with isoflurane (2%), and end-tidal CO_2_, blood O_2_ saturation, electrocardiogram, blood pressure, lung pressure, and body temperature were monitored continuously. A small craniotomy and durotomy were performed over the opercular regions of V1 and a PVC chamber glued to the skull surrounding the craniotomy. On completion of surgery, isoflurane was turned off, anesthesia maintained with sufentanil citrate (6-12 μg/kg/h, i.v.), and paralysis was induced by continuous i.v. infusion of vecuronium bromide (0.3 μg/kg/h) to prevent eye movements. The pupils were dilated with topical atropine, and the corneas were protected with gas-permeable contact lenses. The eyes were refracted using corrective lenses, and the foveae were plotted on a tangent screen using a reverse ophthalmoscope, and periodically remapped throughout the experiment.

#### Electrophysiological recordings

Extracellular recordings (MUA and LFP) were made in parafoveal V1 (4-8° eccentricities) using 24-channel linear electrode arrays (100μm inter-contact spacing, 20μm contact diameter; V-Probe, Plexon, Texas). One penetration was performed using a 32-channel linear probe (100μm spacing; NeuroNexus A32, Michigan). A custom-made guide tube provided mechanical stability for the V-probe recordings. At the beginning of each recording session, the array was positioned normal to the cortical surface under visual guidance using triangulation, and the recording site was stabilized by half filling the chamber with agar; the agar was then covered with silicon oil or saline to prevent it from drying out. The probe was then slowly advanced through the cortical thickness to a depth of 2.0-2.2mm, over a 60-90 minute period, or until LFP signals and spikes could be recorded from the bottom contact through the top third or fourth contact (from the pial surface). At the end of each recording, new craniotomies and durotomies were performed and the recordings targeted to a new cortical site. To facilitate post-mortem visualization of the lesion tracks, the probes were coated with DiI (Molecular Probes, Eugene, OR) prior to insertion.

Data was collected (30kHz sampling rate) and amplified using a 128-channel system (Cerebus,16-bit A-D, Blackrock Microsystems, Salt Lake City, UT). To obtain LFPs, the raw voltage recordings were band-pass filtered (1-100Hz, 2^nd^-order Butterworth filter) and down sampled to 2kHz. MUA was obtained by band-pass filtering (250 Hz-7.5 kHz) the raw signal continuously recorded at a sampling rate of 30 kHz. MUA was thresholded based on signal energy, using the built-in Cerebus program.

#### Receptive Field Mapping and Verticality of the Array

After manually locating the receptive fields (RFs) of neurons across the column, their aggregate minimum response field (mRF) was mapped quantitatively using 0.5° black square stimuli flashed systematically over a 3×3° visual field area (500ms, 5-15 trials, interleaved with 500ms blanks of mean luminance gray). The aggregate spatial mRF of the column was defined as the visual field region in which the patch stimulus evoked a mean response (-2SD of the stimulus evoked response) that was > 2 SD above mean spontaneous activity, and the geometric center of this region was taken as the RF center. All subsequent stimuli were centered on this field. We then determined orientation, eye, spatial and temporal frequency preferences of cells across contacts using 1-1.5° diameter drifting sinusoidal grating patches of 100% contrast presented monocularly. Subsequent stimuli were presented at the optimal parameters for most contacts across the column (unless otherwise specified), and in cases when different contacts showed significantly different stimulus preferences, the experiments were run multiple times using each preferred stimulus. We also measured size tuning across the column using 100% contrast drifting grating patches of increasing size (from 0.1-26°) centered over the aggregate mRF of the column. From these tuning curves we extracted the summation RF (sRF) diameter as the grating diameter at peak response. The latter was later used to create center and annular surround stimuli used to probe surround suppression. To monitor eye movements, the RFs were remapped by hand approximately every 10-20 minutes and stimuli re-centered on the RF if necessary.

For the analysis of sRF sizes shown in **Supplementary Figs. 1A and 2**, size tuning curves were generated by plotting for each stimulus size the mean-2SD of the evoked response (in the first 200 ms) and subtracting from this the mean+2SD of the spontaneous activity; the peak of this curve was chosen as the sRF size.

To ensure that the array was positioned orthogonal to the cortical surface, we used as criteria the vertical alignment of the mapped RF at each contact, and the similarity in the orientation tuning curves across contacts (e.g. **Fig. 2A**). If RFs were misaligned across contacts, the array was retracted and repositioned. Moreover, during offline analysis, we excluded from further analysis all penetrations that were deemed to be non-vertical.

#### Visual Stimuli

Visual stimuli were generated using Matlab (Mathworks Inc, Natick, MA) and presented on a calibrated CRT monitor (Sony, GDM-C520K, 600x800 pixels, 100Hz frame rate, mean luminance 45.7cd/m^2^, at 57cm viewing distance), and their timing was controlled using the ViSaGe system (Cambridge Research Systems, Cambridge, UK). All stimuli were displayed for 500ms, followed by 500 or 750ms interstimulus interval.

To characterize the onset latency of CSD signals evoked by stimuli in the RF and surround, we used two kinds of stimuli: 1. a black square of 0.5° side systematically flashed over a 3x3° visual field areas centered on the columnar RF; and 2. static annular gratings of optimal parameters for the neurons in the column, 2° in width, flashed at distances from the RF center ranging from 0.2° to 6° (measured to the inner edge of the annulus), and presented without a stimulus in the RF.

To characterize the temporal emergence of tuned and untuned surround suppression of spiking responses, we used two kinds of grating stimuli: 1. a 20° diameter drifting grating patch centered on the columnar RF, and presented at the optimal parameters for neurons in the column; and 2. A 2°-width annular static grating of optimal or of orthogonal-to-optimal orientation, flashed at increasing distances from the RF center (from 0.25° outside the edge of the columnar sRF to 6° from the RF center), and presented together with a static grating patch of optimal parameters centered on the columnar RF and matched to the columnar sRF size. The latter stimulus, thus, differed from the former stimulus in that there was always a gap (of the same mean luminance as the grating) between the center and surround gratings. Stimulus presentation was randomized and all grating stimuli were interleaved with presentation of center-only grating patches to activate the RF in isolation.

### Quantification and Statistical Analysis

#### Current Source Density Analysis

Current source density (CSD) was applied to the band-pass filtered (1-100Hz) and trial averaged LFP using the kernel CSD toolbox (Potworowski et al., 2012). CSD was calculated as the second spatial derivative of the LFP signal, which reflects the net local transmembrane currents that generate the LFP.

Specifically, CSD was computed as:

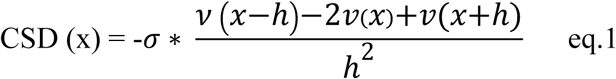

where, *ν* is the voltage (μV), x the point in the extracellular medium at which CSD is calculated, h is the spacing between recording contacts on the linear probe (here 100μm), and *σ* is the connectivity of the cortical tissue (0.4 S/m)(Potworowski et al., 2012). To estimate CSD across layers, we interpolated the CSD every 10*μ*m. The CSD was baseline corrected (Z-scored). In particular, we normalized the CSD of each profile to the standard deviation (SD) of the baseline (defined as 200ms prior to stimulus onset) after subtraction of the baseline mean, as in equation 2:

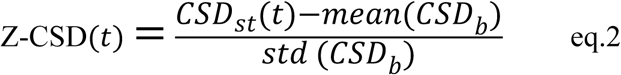

where, *CSD_st_*(*t*) is the computed CSD at each time point (every 0.5ms) after stimulus onset and *CSD_b_* is the computed CSD from 200ms prior to stimulus onset to stimulus onset.

CSD provides information about the current flow in the extracellular medium, and is better suited than LFP to localize input activity (Logothetis et al., 2007). Current sinks (negative voltage deflections, visualized in blue in our CSD maps) in the extracellular medium are thought to reflect integrated subthreshold inputs at postsynaptic dendrites (Mitzdorf, 1985; Mitzdorf and Singer, 1979; Nicholson and Freeman, 1975). We used CSD responses to small stimuli flashed inside the RFs to identify laminar borders (as detailed in the Results). We also used CSD analysis to localize surround-evoked input activity to specific cortical layers. In particular, since surround stimuli presented alone in the absence of direct RF stimulation do not cause significant spiking activity from the recorded cells, the CSD sinks evoked by these stimuli reflect the laminar location of the subthreshold inputs. Measuring the onset latency of these CSD sinks, thus provides us with information about which layers are first activated by surround stimuli.

#### Alignment of Penetrations

To generate average MUAs and CSDs across layers, we aligned the different penetrations using their individual CSDs (as well as the other criteria described in the Results) to identify layers. All our penetrations spanned cortical depths of 1.5-1.6mm, and the bottom of L4C across penetrations was consistently located at depths of 0.9-1mm from the top. This allowed us to align penetrations at the location of the lower border of L4C.

To obtain grand average CSDs (e.g. **Fig. 3D-F**), we half-wave-rectified the individual Z-scored CSD profiles (discarding the positive source values), normalized them between -1 and 0, aligned the individual CSD profiles, and finally averaged the CSD values across aligned penetrations.

#### Latency Analysis of CSD

The onset latency of current sinks in the CSD for the quantitative analysis shown in **Fig. 4** was measured at each interpolated depth as the earliest time bin after stimulus onset in which the CSD amplitude was 5SD below baseline for three consecutive bins. The time of the first bin was taken as the signal onset. For the grand averages shown in **Fig. 3D-F**, onset latency was computed using a SD of 7.

#### Suppression and Orientation Specificity Index

The strength of suppression induced by the surround stimuli was measured as a suppression index (SI), which was computed as:

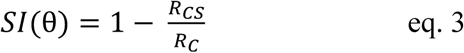

where θ is the surround orientation for the condition, ***R_c_*** is the mean MUA response to the center-only stimulus (in the first 150ms after stimulus onset), and ***R_cs_*** is the mean MUA response to the center+surround stimulus (in the first 150ms). A SI=1 indicates that the surround stimulus completely abolished the response to the center-only stimulus, SI=0 indicates no surround suppression, and SI<1 indicates that the surround stimulus increased the response to the center stimulus alone.

The orientation specificity of surround suppression was measured as an orientation selectivity index (OSI), which was computed as:

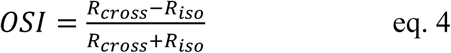

where, ***R_cross_*** and ***R_iso_*** are the response to an iso-oriented and cross-oriented center-surround stimulus, respectively. A OSI=0 indicates no selectivity, while a OSI=1 indicates maximal selectivity.

#### Latency Analysis of Surround Suppression

Peristimulus time histograms (PSTHs) were obtained by convolving a Gaussian filter (10ms bandwidth) with the MUA raster plots, using the chronux toolbox (Buzsaki et al., 2012), providing spikes with 2ms resolution. Then, the mean baseline response was subtracted from the stimulus-evoked response. The onset latency of surround suppression caused by iso-oriented or cross-oriented surround stimuli was estimated to be the time point at which the MUA PSTH in response to the center-surround stimulus diverged significantly from the MUA PSTH in response to the center-only stimulus. This was estimated as follows. First the average MUA response to the center-surround stimulus was subtracted from the average MUA response to the center-only stimulus; second the local minima and maxima of the absolute value of this response difference were found (minima=0 indicates that the two curves intersected); third, the algorithm searched forward in time (up to 300ms after stimulus onset) starting from the first local extrema occurring after the onset of the center-only response, until it found a bin (20ms width) at which the SI reached 0.15 and was followed by a bin with a larger area under the curve (i.e. for which the difference between the two curves was larger). If these criteria held for 3 consecutive bins, then the first bin was chosen as the time of suppression onset. Essentially, based on these constraints, the time of suppression could only be equal or larger than the response onset to the center-only stimulus. The response onset to the center-only stimulus was taken to be the time point at which the PSTH reached 10% of maximum value (Bokil et al., 2010). For the boxplots of **Fig. 6A,B,D** this analysis was performed only on MUA responses that showed an SI≥0.15 (measured over 150ms after stimulus onset, because after this time window the response was typically back to baseline).

The onset latency of tuned surround suppression was measured, using the same approach as described above, as the time point at which the PSTHs in response to iso-oriented and cross-oriented center-surround stimuli diverged significantly (i.e. the absolute OSI value was >0.1 for 3 consecutive bins). An additional constraint, here, was that the onset of tuned suppression could not precede in time the onset of untuned suppression (i.e. the onset of suppression caused by either an iso-oriented or a cross-oriented center-surround stimulus, whichever was faster). For the boxplots of **Fig. 6C** this analysis was performed only on MUA responses that showed an SI≥0.15 for either iso- or cross-oriented center-surround stimuli (over the 150ms after stimulus onset).

We also compared our method to one that computes the latency by taking into account the trial variability, using an approach similar to that used in Henry et al (Henry et al., 2013); this method compares the cumulative spike counts during stimulus presentation in the center-only vs. the center-surround stimuli. The cumulative spike counts were generated by bootstrap resampling (with replacement, 5000 iteration) from the population of trials at each time bin (10ms width). The time bin was chosen as the time of onset latency, if the spike count in response to the center-only stimulus was larger than the spike count in response to the center-surround stimulus for at least 95% of the time. We found this method was only effective to measure latency under conditions that evoked strong surround suppression or in which surround suppression was sharply tuned (as also discussed in Henry et al. (2013). However, with this method the latency estimate increases as the change in response is scaled down, i.e. for weaker surround suppression or more weakly tuned suppression, as is the case for far surround suppression. We do not report the results of this analysis, as we found it to be ineffective to measure onset latency for far surround suppression.

#### Histology and Track Reconstruction

On completion of the recording session, the animal was perfused transcardially with 4% paraformaldehyde in 0.1M phosphate buffer. The occipital pole was frozen-sectioned at 40μm sagittally. DiI-labeled tracks were visualized under fluorescence to ascertain verticality of the array and verify cortical layer assignment (e.g. **Fig. 2B**). Adjacent tissue sections were counterstained for cytochrome oxidase for identifying cortical layers as well as the location of the electrode track (visible as a discoloration in staining).

